# High burden of *Mycoplasma genitalium* and other reproductive tract infections among pregnant women in Papua New Guinea

**DOI:** 10.1101/2020.05.22.109983

**Authors:** Michelle J. L. Scoullar, Philippe Boeuf, Elizabeth Peach, Ruth Fidelis, Kerryanne Tokmun, Pele Melepia, Arthur Elijah, Catriona S. Bradshaw, Glenda Fehler, Peter M. Siba, Simon Erskine, Elisa Mokany, Elissa Kennedy, Alexandra J. Umbers, Stanley Luchters, Leanne J. Robinson, Nicholas C. Wong, Andrew Vallely, Steven G. Badman, Lisa M. Vallely, HMHB Study Team, Freya J. I. Fowkes, Christopher Morgan, William Pomat, Brendan S. Crabb, James G. Beeson

## Abstract

There is a pressing need for detailed knowledge of the range of pathogens, extent of co-infection and clinical impact of reproductive tract infections (RTIs) among pregnant women. Here, we report on RTIs (*Mycoplasma genitalium, Chlamydia trachomatis*, *Neisseria gonorrhoeae*, *Trichomonas vaginalis*, *Treponema pallidum subspecies pallidum,* bacterial vaginosis and vulvovaginal candidiasis) and other sexual and reproductive health indicators among 699 pregnant women in Papua New Guinea (PNG). We found widespread *M. genitalium* infection (12.5% of women), the first time this pathogen has been reported in PNG, with no evidence of macrolide resistance. Most pregnant women (76.2%) had at least one RTI, most of which are treatable. Excluding syphilis, sexually-transmitted infections were detected in 37.8% women. Syndromic management of infections is greatly inadequate and there was remarkably little use of contraception; 98.4% report never having used barrier contraception. This work has implications for improving maternal and child health in PNG.

**ARTICLE SUMMARY LINE:** This first report of *Mycoplasma genitalium* in Papua New Guinea finds a high burden (12.5%) among 699 pregnant women. Additionally, more than one in two women were positive for a treatable reproductive tract infection associated with poor health outcomes.

## BACKGROUND

Reproductive Tract Infections (RTIs), including Sexually Transmitted Infections (STIs), are a preventable health burden. Of over 30 well recognised STIs, four are referred to as curable STIs: *Chlamydia trachomatis*, *Neisseria gonorrhoeae*, *Trichomonas vaginalis* and *Treponema pallidum subspecies pallidum* (syphilis). An estimated 376.4 million new adult cases of these four infections occur annually and the World Health Organization (WHO) Western Pacific Region has the highest number of annual incident cases, estimated at 142 million(1–3). Other RTIs, such as bacterial vaginosis (BV) and *Candida* spp. causing vulvovaginal candidiasis (VVC) are common, but global estimates less available and challenged by differing diagnostic methodologies for BV(4) and that *Candida* spp can be pathogenic and commensal. Notwithstanding this, BV estimates range from 8% to 51% in pregnant women(5) and *Candida* spp. can be isolated from vaginal samples in 20-30% of asymptomatic women and 40% of symptomatic women (6). Women with RTIs can experience substantial pain and discomfort and may be subject to debilitating stigma(7). Complications can include pelvic inflammatory disease (PID), infertility, increased risk of acquisition and transmission of other STIs; and in pregnancy can result in miscarriage, stillbirth, preterm birth (PTB), neonatal death, and serious neonatal morbidities such as blindness, congenital malformations and lifelong disability(1, 8, 9).

The curable STI *Mycoplasma genitalium* has recently emerged as an important cause of poor sexual health, associated with PID, cervicitis, miscarriage and PTB(10, 11). Data on prevalence is more limited than for other curable STIs, available reports range from below 1% in the general adult population to 15.9% in high risk groups(12, 13), and in pregnancy from 0.7% in the United Kingdom(14) to 11.9% in the Solomon Islands(15). Importantly, there has been a marked decline in azithromycin cure over the past decade, with some areas reporting macrolide resistance as high as 68%(16–18). However, in many regions, the importance of *M. genitalium* is largely unrecognised due to a lack of data on prevalence and drug sensitivity.

Papua New Guinea (PNG) is a Pacific nation with more than 8.5 million people(19) and high rates of *C. trachomatis*, *N. gonorrhoeae* and *T. vaginalis* that exceed those seen in other high burden regions such as sub-Saharan Africa (1, 20). However, there are no reports on the prevalence of *M. genitalium*. In this study we evaluated the prevalence of *M. genitalium* and molecular markers of resistance, as well as rates of other RTIs and curable STIs, among pregnant women attending antenatal clinics in East New Britain Province (ENBP) of PNG. We investigated the relationships between different RTIs, factors associated with infection, and compared the performance of syndromic management to more accurate molecular and rapid test diagnostics.

## METHODS

### Study site and population

This study uses cross-sectional baseline data from 699 pregnant women attending their first antenatal clinic (ANC1) who were enrolled into a prospective cohort study “Healthy Mothers Healthy Babies” (HMHB) undertaken across five health facilities in ENBP, PNG. Enrolment occurred between March 2015 and June 2017. Women aged 16 years or older, living in the facilities’ catchment area and attending clinic for the first time in the current pregnancy regardless of gestation, were eligible to participate. At each site women were randomly selected and invited to participate. After eligibility screening and informed consent, a questionnaire was administered by a trained research officer. We collected socio-demographic and clinical information, after which biological samples were obtained, including self-collected vaginal swabs, urine, capillary finger prick and venous blood samples. All abnormal results available at the point-of-care (urine dipstick, syphilis, malaria, haemoglobin) were communicated directly to the participant, the health care provider and documented in the individual client-held health record book, at the time of interview, to enable treatment by the health care provider.

### Study procedures

Routine antenatal care was provided by the health facility staff in line with PNG national guidelines (Supplemental methods)(21, 22). As per national STI syndromic management guidelines, women reporting current abnormal vaginal discharge receive: intravaginal nystatin pessaries (100,000 units, twice daily for 7 days) or a clotrimazole pessary once followed by intravaginal clotrimazole cream for 7 days; amoxycillin 2g orally (PO); probenecid 1g PO; augmentin 2 tablets PO, and azithromycin 1g PO.

Two self-collected vaginal swabs were provided by each participant; one was used to prepare a vaginal smear for microscopy, and each swab was then placed into a DNA preserving transport medium tube, stored in a chilled cooler and returned to the laboratory at the end of each day. Urine samples were collected in a sterile container, placed on ice and later stored at −20 degrees Celsius. Swabs were tested in ENB for *C. trachomatis, N. gonorrhoea* and *T. vaginalis;* other samples were shipped to Burnet Institute Melbourne, Australia in batches. In cases where testing was delayed, specimens were stored at 2-7°C or at −20°C if an extended delay was anticipated. Molecular testing usually occurred within seven days with results communicated back to participants and treatment advised for themselves and their partner.

### Laboratory methods

The GeneXpert molecular platform (Cepheid, Sunnyvale CA, USA) was used to test vaginal and urine specimens at the Burnet / PNG Institute of Medical Research (PNG IMR) laboratory, Vunapope Hospital Kokopo for *C. trachomatis, N. gonorrhoea and T. vaginalis*. Lifetime exposure to *T. pallidum* (syphilis) was determined to be positive if women tested positive by Alere Determine™ Syphilis TP (Abbott, Illinois, USA). This was initially performed at health facilities in line with national guidelines; however, inconsistency in test availability at different sites meant the research team supplied rapid tests for study women.

Shipped vaginal specimens were stored at the Burnet Institute Melbourne at −20 or −80°C until all samples had been received. After thawing, genomic DNA was extracted using the QIAamp^®^ BiOstic® Bacteremia DNA kit (QIAGEN N.V, Venlo, The Netherlands). To optimise this protocol for vaginal specimens rather than cultured blood, the total volume of sample required was adjusted to 450μl. Extracted DNA was tested for *M. genitalium* using the *ResistancePlus*® MG kit (SpeeDx, Sydney, Australia) which uses the PlexPCR® and PlexPrime® technologies (SpeeDx) for concurrent amplification of *M. genitalium* and detection of five *M. genitalium* point mutations (A2058G, A2058C, A2058T, A2059G, and A2059C) within the macrolide resistance-determining region (MRDR) of the 23S rRNA gene(17). The *ResistancePlus*® MG kit is commercially available, approved by Therapeutic Goods Administration Australia, and has a sensitivity and specificity of 98% and 100% for *M. genitalium* and 96% and 100% for macrolide resistance mutations(23–25). Gram-stained vaginal smears were read by an experienced microscopist at Melbourne Sexual Health Centre. BV diagnosis was based on Nugent scoring (BV, Nugent score 7 to 10) and presumptive diagnosis of candidiasis was based on observation of pseudohyphae and or budding yeasts. Because the diagnosis of *M. genitalium,* BV and VVC was on stored samples many months after collection it was not possible to specifically treat those women who tested positive beyond any syndromic management they would have received as part of national guidelines.

### Data management and statistical analysis

Questionnaire responses were entered by research officers directly into an electronic tablet (e-tablet) using a study specific questionnaire on the platform Mobile Data Studio 7.3 (MDS, CreativityCorp Pty Ltd, Perth, Australia), and stringent data management protocols were in place (supplemental methods).

Exposures of interest included clinic details (enrolment clinic; rural or urban); participant characteristics at enrolment and relevant obstetric history (see supplemental methods for a full list of variables and definitions). Outcomes measured were *M. genitalium, C. trachomatis, N.gonorrhoea, T. vaginalis, T. pallidum* (syphilis), BV and VVC. The association between exposures and *C. trachomatis, N. gonorrhoea, T. vaginalis, M. genitalium* and syphilis were assessed using logistic regression. All variables of interest were included in univariable analyses and variables associated with the outcome at p<0.10 in univariable analyses were included in a multivariable model along with the pre-specified variable of enrolment clinic site.

### Ethical considerations and informed consent procedures

All participants provided individual written, informed consent. Ethical approval was provided from the Medical Research Advisory Committee of the PNG National Department of Health (No. 14.27), the PNG IMR Institutional Review Board (No. 1114) and the Human Research Ethics Committee of the Alfred Hospital (No. 348/14) in Australia. Provincial approval was obtained from the East New Britain Province (ENBP) Executive Committee, participating facilities and a series of community engagement meetings provided broader community support and assent for the study.

## RESULTS

A total of 699 pregnant women were enrolled at five antenatal clinics in East New Britain Province (ENBP). Median maternal age was 26 years (interquartile range, IQR 22-30), a quarter of women were primigravida (177/699, 25.4%), the majority were married / co-habiting (663/697, 95.1%) and 46.5% had only completed primary school (325/698) (Table 1).

**TABLE 1:**
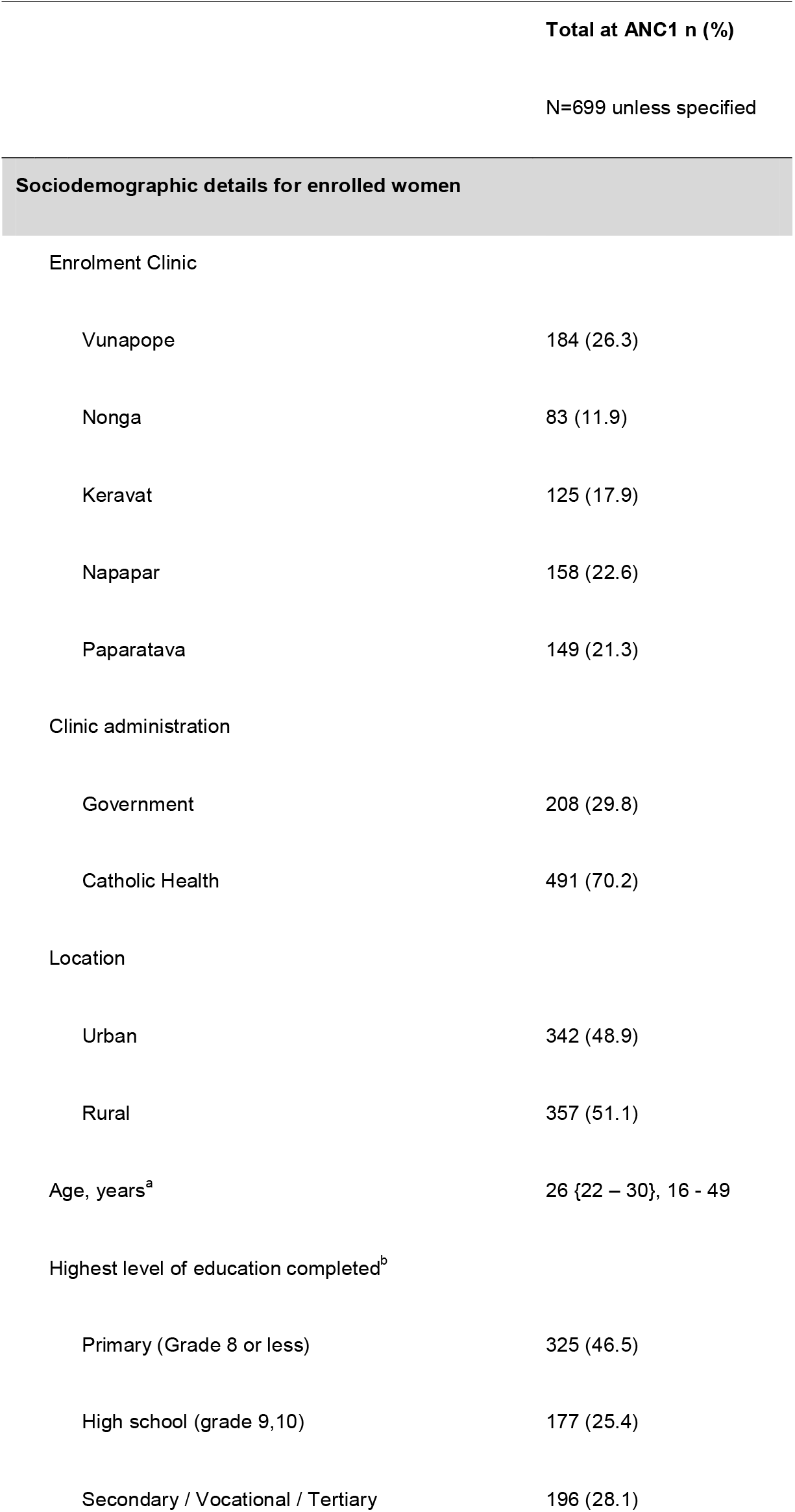
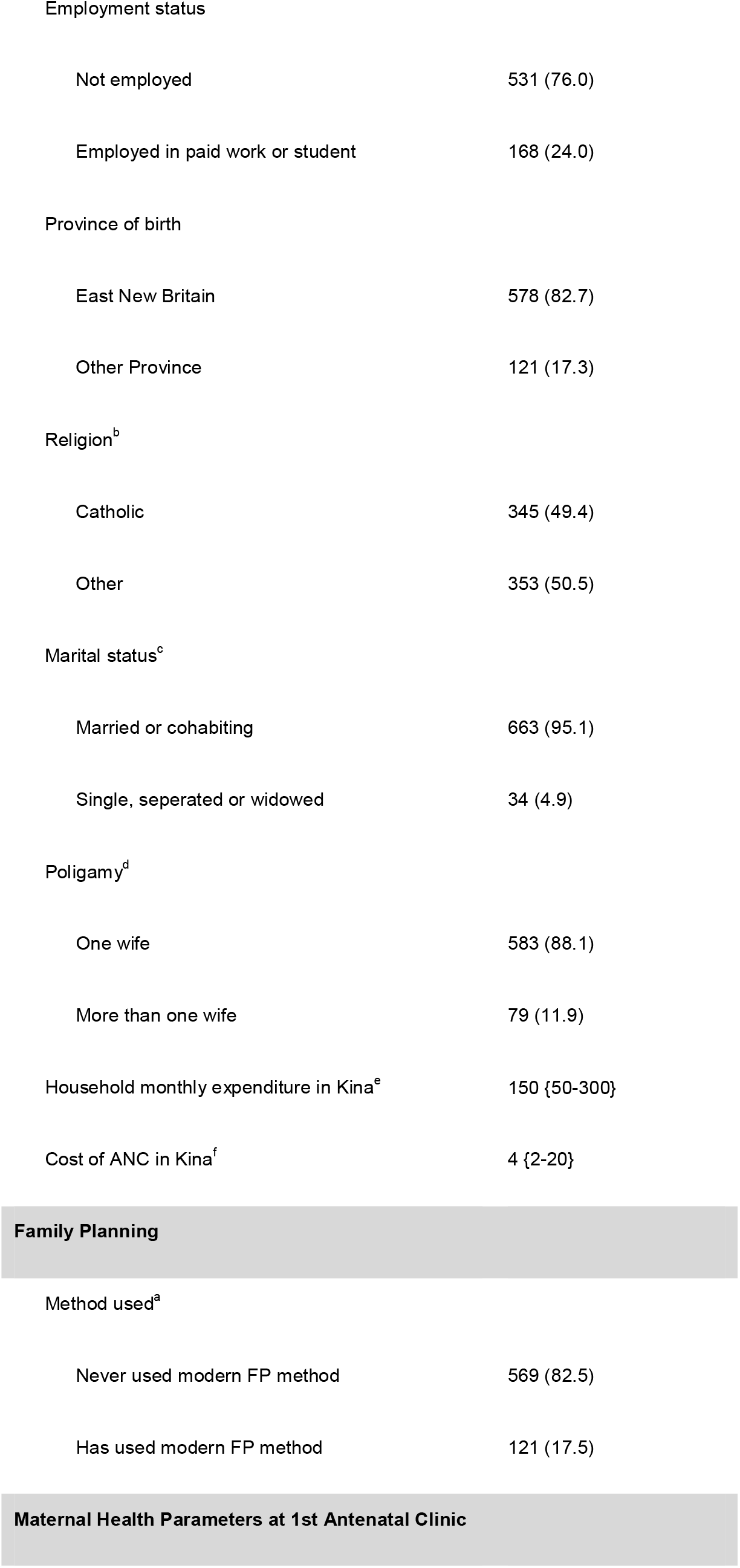
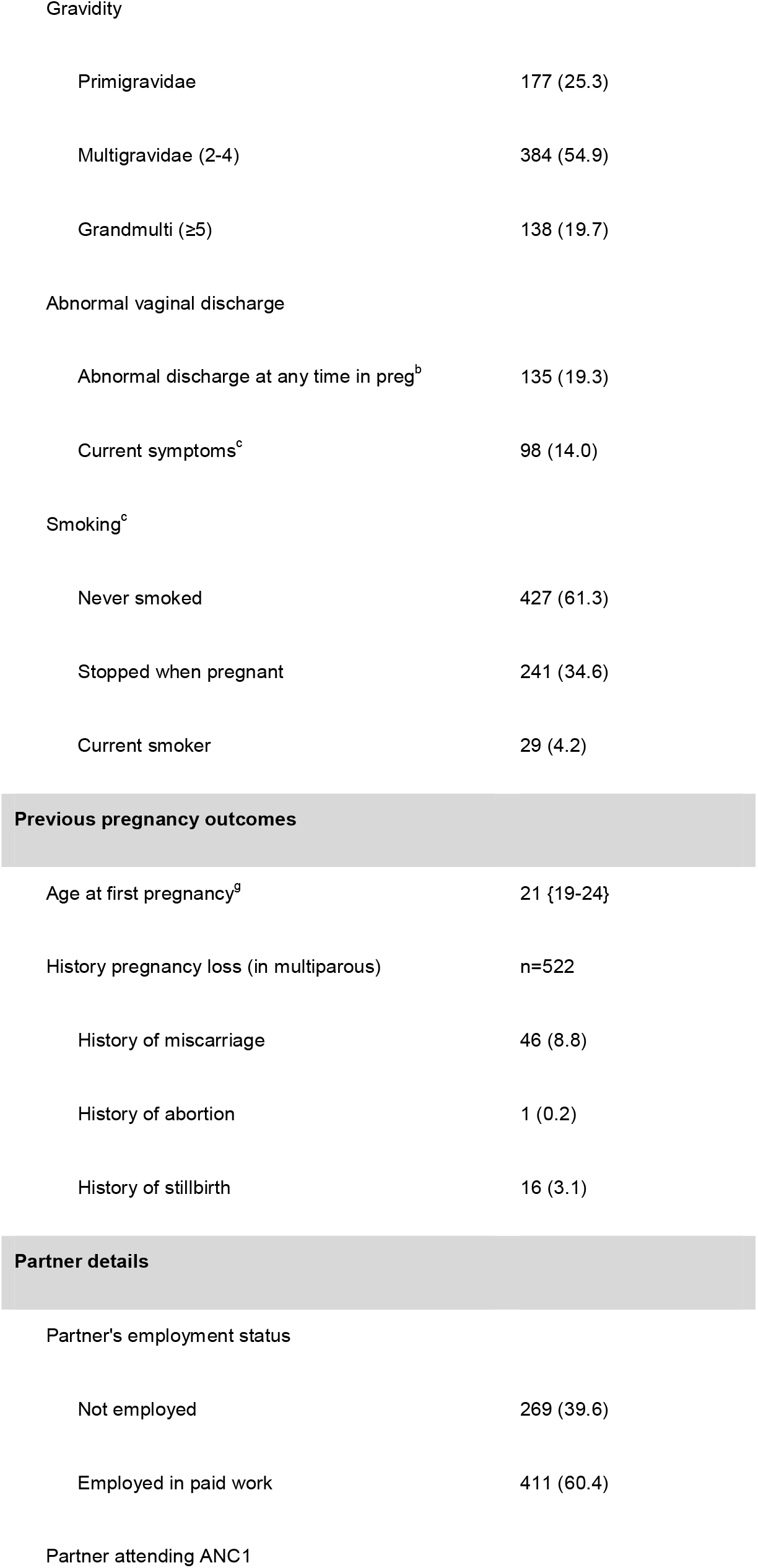
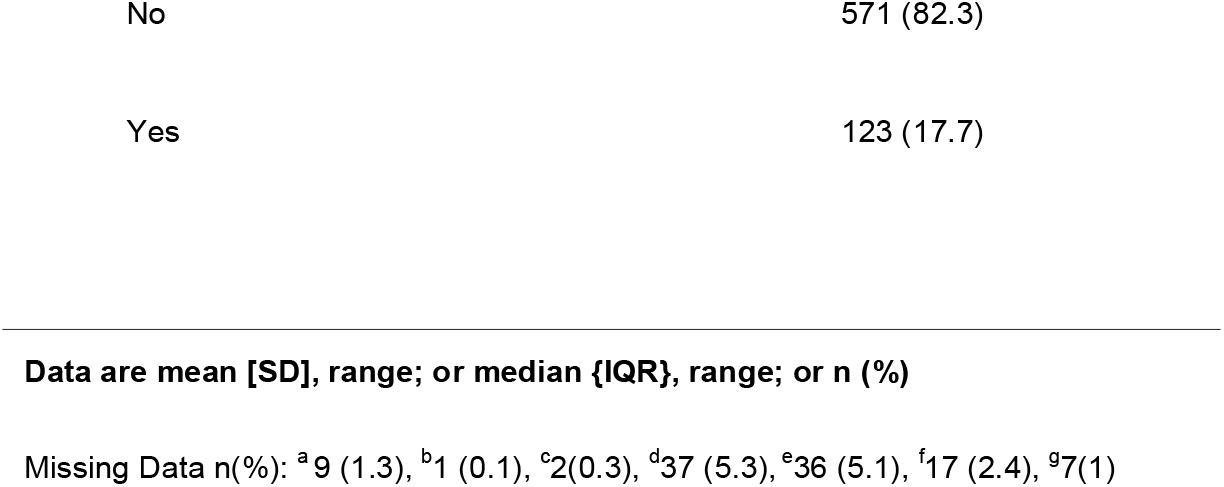
Socio-demographic characteristics and obstetric history of women at first antenatal clinic visit in East New Britain, PNG.

Current abnormal vaginal discharge was reported by 14.0% (98/697), an additional 37 women reported abnormal vaginal discharge at some point in this pregnancy prior to clinic attendance (total symptomatic discharge in pregnancy 19.3%, 135/698). Most women (82.5%, 569/690) had never used a modern method of contraception, with only 11 women (1.6%, 11/690) reporting having ever used either a male or female condom.

### High burden of reproductive tract infections in pregnancy

Of the 699 women enrolled, prevalence of *M. genitalium* was 12.5% (78/625; 95% CI 10.0-15.4), with no evidence of macrolide resistant mutations (Table 2). Other RTIs were: *C. trachomatis* 19.1% (122/640; 95% CI 16.1-22.4), *N. gonorrhoeae* 5.5% (35/640; 95% CI 3.9-7.6), *T. vaginalis* 20.2% (117/581; 95% CI 17.0-23.7) and prevalence of lifetime exposure to syphilis was extremely high at 18.1% (79/437; 95% CI 14.6-22.1). Of the 503 vaginal smears available for microscopy, BV prevalence was 25.7% (129/503; 95% CI 21.9-29.7) and 39.4% (198/503; 95% CI 35.1-43.8) had presumptive candidiasis detected on microscopy (VVC).

**TABLE 2.** Prevalence of reproductive tract infections among pregnant women in ENB PNG.

| Reproductive Tract Infection | Obs | Freq | Prevalence % (95% CI) |
| --- | --- | --- | --- |
| *Mycoplasma genitalium* | 625 | 78 | 12.5 (10.0 - 15.4) |
| *Chlamydia trachomatis* | 640 | 122 | 19.1 (16.1 - 22.4) |
| *Neisseria gonorrhoeae* | 640 | 35 | 5.5 (3.9 - 7.6) |
| *Trichomonas vaginalis* | 581 | 117 | 20.2 (17.0 - 23.7) |
| Syphilis (Alere Determine) | 437 | 79 | 18.1 (14.6 - 22.1) |
| Bacterial Vaginosis | 503 | 129 | 25.7 (21.9 - 29.7) |
| Vulvovaginal candidiasis | 503 | 198 | 39.4 (35.1 - 43.8) |
| **At least 1 of a group of RTIs<sup>a</sup>** |  |  |  |
| At least 1 current RTI<sup>a</sup> | 357 | 272 | 76.2 (71.5 - 80.6) |
| At least 1 current STI<sup>b</sup> | 485 | 183 | 37.8 (33.5 - 42.3) |
| At least 1 of MG, CT, NG, TV or Syphilis | 302 | 144 | 47.7 (42.0 - 53.5) |
| At least 1 of MG, CT, NG, TV or BV | 357 | 197 | 55.2 (49.9 - 60.5) |
| At least 1 GeneXpert diagnosed infection<sup>c</sup> | 546 | 175 | 32.1 (28.2 - 36.2) |
| At least 1 vaginal infection<sup>d</sup> | 409 | 281 | 68.8 (64.0 - 73.2) |
| At least 1 of BV or VVC | 503 | 293 | 58.3 (53.9 - 62.6) |
| **Multiple current STIs** |  |  |  |
| Any 2 current STIs | 661 | 75 | 11.4 (9.1 - 14.1) |
| Any 3 current STIs | 536 | 15 | 2.8 (1.6 - 4.6) |

<sup>a</sup>Participants result included only if they had all tests done for each of the infections within group of RTIs
<sup>a</sup> current RTI includes at least 1 of MG, CT, NG, TV, BC or VVC (syphilis not included)
<sup>b</sup> current STI includes at least 1 of MG, CT, NG, TV (syphilis not included)
<sup>c</sup> GeneXpert diagnosed infections include any 1 of CT, NG or TV
<sup>d</sup> vaginal infections includes at least 1 of BV, TV or VVC
RTI (reproductive tract infection), STI (Sexually transmitted infection), MG (*Mycoplasma genitalium*), CT (*Chlamydia trachomatis*), NG (*Neisseria gonorrhoea*), TV (*Trichomonas vaginalis*), BV (Bacterial Vaginosis), VVC (Vulvovaginal candidiasis)

The majority of women (76.2%, 272/357), had at least one current RTI (BV, VVC, *M. genitalium, C. trachomatis, N. gonorrhoea* or *T. vaginalis)*, with at least one current STI present in 37.8% (183/485, *M. genitalium, C. trachomatis, N. gonorrhoea* or *T. vaginalis*). One in three women (32.1%, 175/546) had a current STI diagnosed using GeneXpert (*C. trachomatis, N. gonorrhoea* or *T. vaginalis)*, 11.4% (75/661) had at least two co-existing STIs, 15 women (2.8%, 15/536) had at least three infections and one woman had all four STIs.

### Relationships between infections

Of the 78 women with *M. genitalium*, 28 women (35.9%) had a concurrent STI detected: 20 (25.6%) had co-infection with *C. trachomatis*, 13 (16.7%) had co-infection with *T. vaginalis* and 6 (7.7%) had *N. gonorrhoeae* (Figure 1 and Table S2). Co-infections were most frequent in women positive for *N. gonorrhoea* (80%, 28/35), with the majority of these women having a co-infection with *C. trachomatis* (71.4%, 25/35) followed by *T. vaginalis* (22.8%, 8/35) and *M. genitalium* (17.1%, 6/35). Syphilis was not included in estimates of co-infections as it was not known if their exposure represented current or previous infection.

**FIGURE 1:**
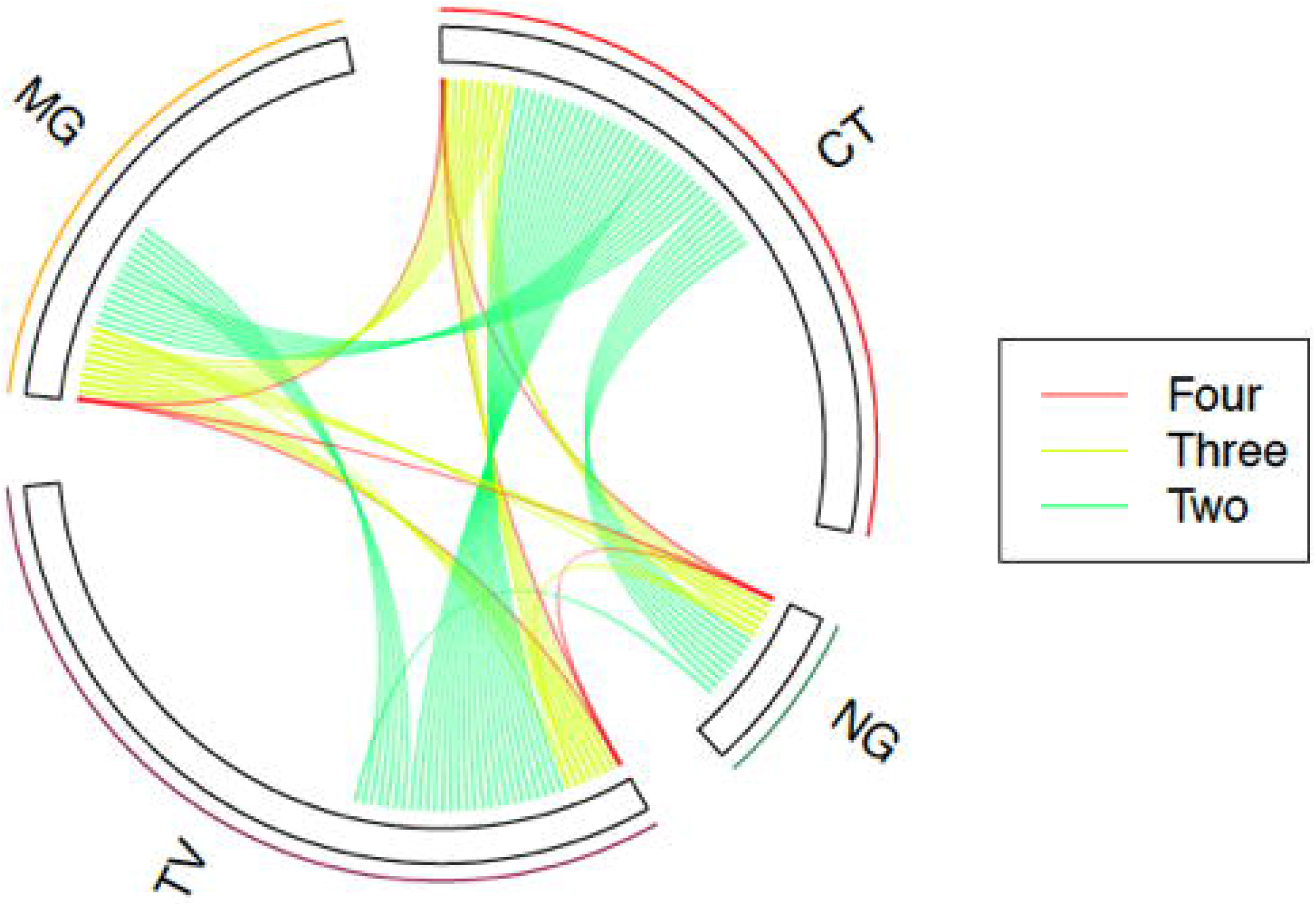
Relationships between current STIs: *Mycoplasma genitalium* (MG)*, Chlamydia trachomatis* (CT), *Neisseria gonorrhoea* (NG) *and Trichomonas vaginalis* (TV). Each line connecting two or more infections represents one participant. Mono infections are represented by the space under each STI with no lines.

Of 129 women with bacterial vaginosis, 37.2% (48/129) had a co-infection, the most common of which was *C. trachomatis* 22.5% (29/129), followed by *T. vaginalis* (12.4%, 16/129), *M. genitalium* (11.6%, 15/129) and *N. gonorrhoea* (7.7%, 10/129) (Table S3).

### Relationship between abnormal vaginal discharge and infection

We compared clinical symptoms used in PNG national treatment guidelines (current abnormal vaginal discharge) with an alternative question of abnormal vaginal discharge currently or at any time in pregnancy up to their first antenatal clinic visit (ANC1) (Table 3). A total of 98 women (14.1%, 98/697) had current symptoms that would have prompted treatment as per PNG national guidelines. An additional 37 women experienced abnormal vaginal discharge earlier in the pregnancy, but no current symptoms, and therefore would not normally receive treatment.

**TABLE 3:** Number of women with a Reproductive Tract Infection and symptoms (abnormal vaginal discharge)

|  | Question as per syndromic management |  |  | Alternative question |  |  |
| --- | --- | --- | --- | --- | --- | --- |
|  | Do you currently have any abnormal vaginal discharge? |  |  | Have you experienced any abnormal vaginal discharge earlier in the pregnancy or now?^ |  |  |
|  | No n(%) | Yes n(%) | Total | No n(%) | Yes n(%) | Total |
|  | 599 (85.9) | 98 (14.1) | 697 | 563 (80.7) | 135 (19.3) | 698 |
| **Reproductive Tract Infection** |  |  |  |  |  |  |
| *Mycoplasma genitalium* | 66 (84.6) | 12 (15.4) | 78 | 61 (78.2) | 17 (21.8) | 78 |
| *Chlamydia trachomatis* | 98 (80.3) | 24 (19.7) | 122 | 90 (73.8) | 32 (26.2) | 122 |
| *Neisseria gonorrhoeae* | 28 (80.0) | 7 (20.0) | 35 | 26 (74.3) | 9 (25.7) | 35 |
| *Trichomonas vaginalis* | 94 (80.3) | 23 (19.7) | 117 | 83 (70.9) | 34 (29.1) | 117 |
| Syphilis (Alere Determine) | 68 (86.1) | 11 (13.9) | 79 | 65 (82.3) | 14 (17.7) | 79 |
| Bacterial Vaginosis | 109 (84.5) | 20 (15.5) | 129 | 101 (78.3) | 28 (21.7) | 129 |
| Vulvovaginal candidiasis | 160 (80.8) | 38 (19.2) | 198 | 146 (73.7) | 52 (26.3) | 198 |
| **At least 1 of a group of RTIs** |  |  |  |  |  |  |
| At least 1 current RTI<sup>a</sup> | 224 (82.3) | 48 (17.6) | 272 | 204 (75.0) | 68 (25.0) | 272 |
| At least 1 current STI<sup>b</sup> | 154 (84.1) | 29 (15.8) | 183 | 141 (77.0) | 42 (22.9) | 183 |
| At least 1 GeneXpert diagnosed infection<sup>c</sup> | 141 (80.6) | 34 (19.4) | 175 | 127 (72.6) | 48 (27.4) | 175 |
| At least 1 vaginal infection<sup>d</sup> | 228 (81.1) | 53 (18.9) | 281 | 207 (73.7) | 74 (26.3) | 281 |
| At least 1 of BV or VVC | 242 (82.6) | 51 (17.4) | 293 | 222 (75.8) | 71 (24.2) | 293 |

| Multiple current STIs |  |  |  |  |  |  |
| --- | --- | --- | --- | --- | --- | --- |
| Any 2 current STIs | 58 (77.3) | 17 (22.7) | 75 | 53 (70.7) | 22 (29.3) | 75 |

<sup>a</sup> 1 woman who responded yes to the second question had a missing response to the first question
<sup>a</sup> active RTI includes at least 1 of MG, CT, NG, TV, BC or VVC (syphilis not included)
<sup>b</sup> active STI includes at least 1 of MG, CT, NG, TV (syphilis not included)
<sup>c</sup> GeneXpert diagnosed infections include any 1 of CT, NG or TV
<sup>d</sup> vaginal infections includes at least 1 of BV, TV or VVC
MG (Mycoplasma genitalium), CT (Chlamydia trachomatis), NG (Neisseria gonorrhoea), TV (Trichomonas vaginalis), BV (Bacterial Vaginosis), VVC (Vulvovaginal candidiasis)

Most current STIs were asymptomatic. Neither the standard question used in national guidelines nor the alternative question performed well as a marker of infection. Of those women with an STI detected, no current symptoms were reported by 84.1% (154/183), and even with the more inclusive alternative question, 77.0% (141/183) reported no history of symptoms either currently or at any time earlier in the pregnancy. Of the women with *M. genitalium*, 78.2% (61/78) had no symptoms either currently or at any time in the pregnancy up to ANC1, and only 12 women (15.4%, 12/78) would have been treated using current syndromic management.

Asking if women had any symptoms in pregnancy up to and including the current time (the alternative question) was consistently more sensitive for any individual infection or group of infections compared the standard question (Figure 2); despite this, sensitivity remained below 30%. The alternative question did perform better to distinguish those women with *T. vaginalis* infection (p=0.005) (Table S4), and VVC (p=0.007) compared to the current question.

**FIGURE 2:**
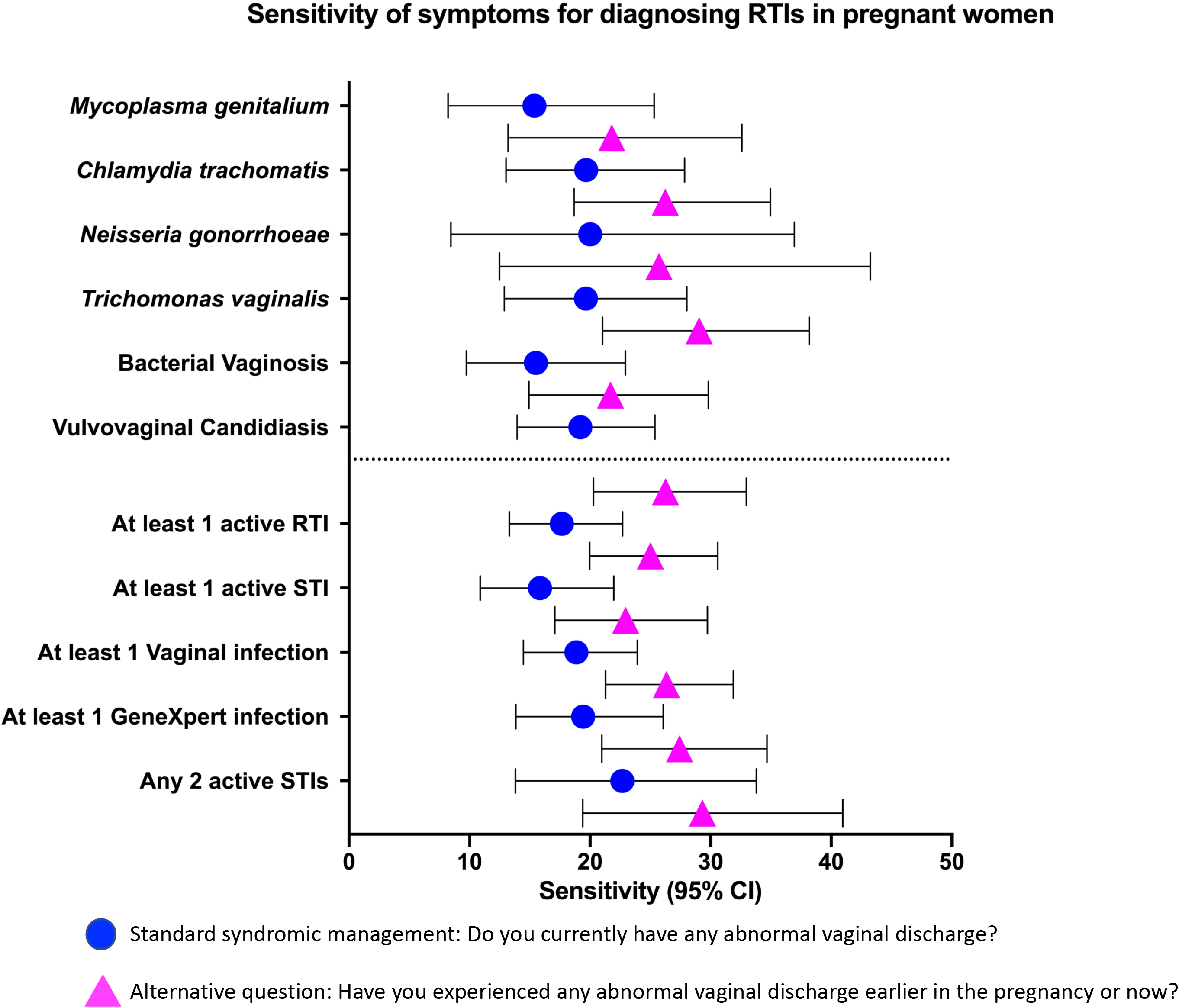
Sensitivity of syndromic management using the standard question as per PNG national guidelines “Do you currently have any abnormal vaginal discharge?”, or an alternative question “Have you experienced any abnormal vaginal discharge earlier in the pregnancy or now?”.

### Factors associated with curable STI infection

There was no factor in either univariable (Table S5) or multivariable (Table 4) analysis found to be associated with an increased odds of *M. genitalium* infection. Given first pregnancy and younger women had a collinear relationship, only pregnancy number was included in the multivariable analysis. Those in their first pregnancy, employment, those who were single or separated and those with abnormal vaginal discharge at any time in the pregnancy up to and including ANC1 were all identified as risk factors for different STIs to varying degrees of statistical significance. Not having used a modern method of contraception appeared to be an important risk factor for a number of STIs in univariable analyses; however this association weakened in multivariable analysis. Risk of exposure to syphilis appeared to vary depending on clinic site.

**TABLE 4:**
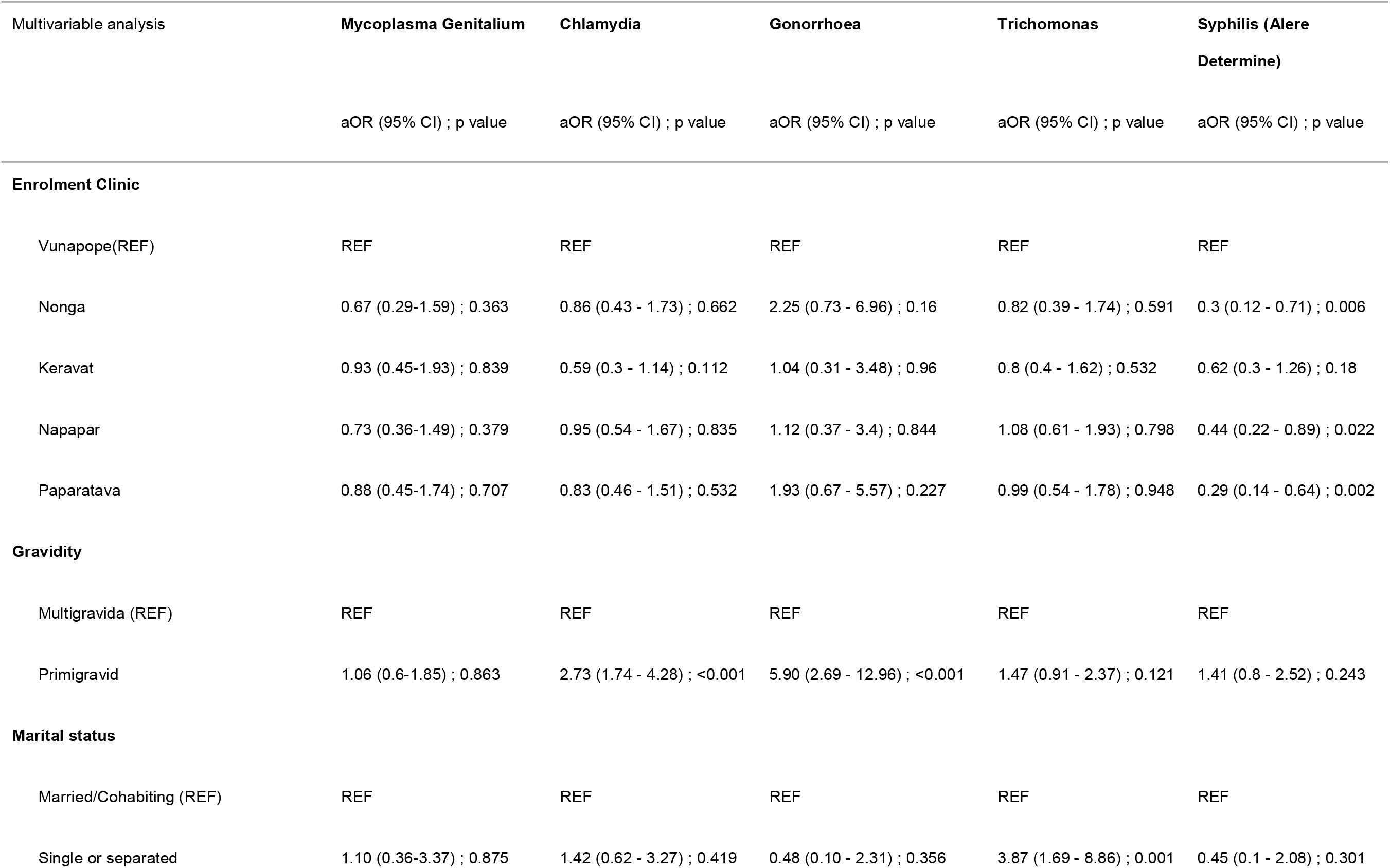
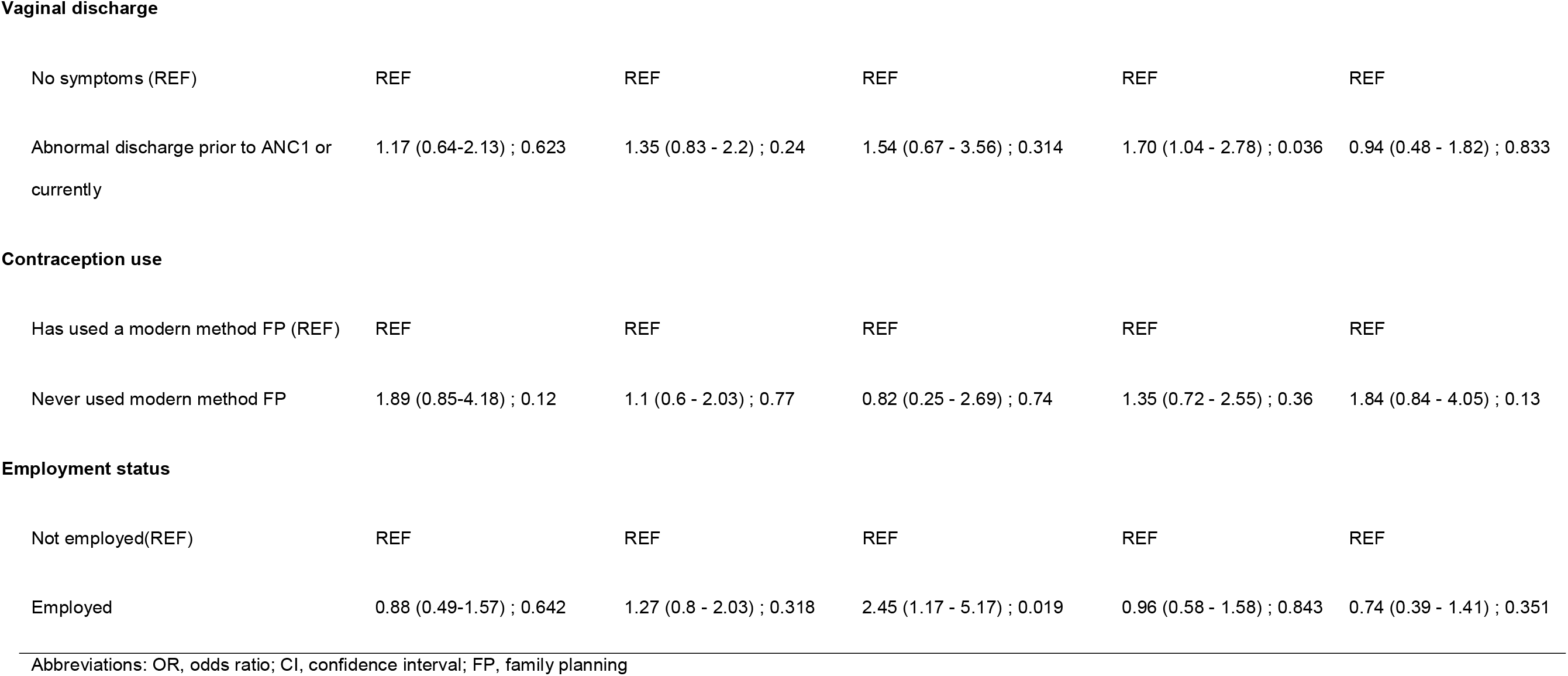
Multivariable analysis of factors associated with curable Sexually Transmitted Infections.

| Multivariable analysis | **Mycoplasma Genitalium** | **Chlamydia** | **Gonorrhoea** | **Trichomonas** | **Syphilis (Alere Determine)** |
| --- | --- | --- | --- | --- | --- |
|  | aOR (95% CI) ; p value | aOR (95% CI) ; p value | aOR (95% CI) ; p value | aOR (95% CI) ; p value | aOR (95% CI) ; p value |
| **Enrolment Clinic** |  |  |  |  |  |
| Vunapope(REF) | REF | REF | REF | REF | REF |
| Nonga | 0.67 (0.29-1.59) ; 0.363 | 0.86 (0.43 - 1.73) ; 0.662 | 2.25 (0.73 - 6.96) ; 0.16 | 0.82 (0.39 - 1.74) ; 0.591 | 0.3 (0.12 - 0.71) ; 0.006 |
| Keravat | 0.93 (0.45-1.93) ; 0.839 | 0.59 (0.3 - 1.14) ; 0.112 | 1.04 (0.31 - 3.48) ; 0.96 | 0.8 (0.4 - 1.62) ; 0.532 | 0.62 (0.3 - 1.26) ; 0.18 |
| Napapar | 0.73 (0.36-1.49) ; 0.379 | 0.95 (0.54 - 1.67) ; 0.835 | 1.12 (0.37 - 3.4) ; 0.844 | 1.08 (0.61 - 1.93) ; 0.798 | 0.44 (0.22 - 0.89) ; 0.022 |
| Paparatava | 0.88 (0.45-1.74) ; 0.707 | 0.83 (0.46 - 1.51) ; 0.532 | 1.93 (0.67 - 5.57) ; 0.227 | 0.99 (0.54 - 1.78) ; 0.948 | 0.29 (0.14 - 0.64) ; 0.002 |
| **Gravidity** |  |  |  |  |  |
| Multigravida (REF) | REF | REF | REF | REF | REF |
| Primigravid | 1.06 (0.6-1.85) ; 0.863 | 2.73 (1.74 - 4.28) ; <0.001 | 5.90 (2.69 - 12.96) ; <0.001 | 1.47 (0.91 - 2.37) ; 0.121 | 1.41 (0.8 - 2.52) ; 0.243 |
| **Marital status** |  |  |  |  |  |
| Married/Cohabiting (REF) | REF | REF | REF | REF | REF |
| Single or separated | 1.10 (0.36-3.37) ; 0.875 | 1.42 (0.62 - 3.27) ; 0.419 | 0.48 (0.10 - 2.31) ; 0.356 | 3.87 (1.69 - 8.86) ; 0.001 | 0.45 (0.1 - 2.08) ; 0.301 |
| **Vaginal discharge** |  |  |  |  |  |
| No symptoms (REF) | REF | REF | REF | REF | REF |
| Abnormal discharge prior to ANC1 or currently | 1.17 (0.64-2.13) ; 0.623 | 1.35 (0.83 - 2.2) ; 0.24 | 1.54 (0.67 - 3.56) ; 0.314 | 1.70 (1.04 - 2.78) ; 0.036 | 0.94 (0.48 - 1.82) ; 0.833 |
| **Contraception use** |  |  |  |  |  |
| Has used a modern method FP (REF) | REF | REF | REF | REF | REF |
| Never used modern method FP | 1.89 (0.85-4.18) ; 0.12 | 1.1 (0.6 - 2.03) ; 0.77 | 0.82 (0.25 - 2.69) ; 0.74 | 1.35 (0.72 - 2.55) ; 0.36 | 1.84 (0.84 - 4.05) ; 0.13 |
| **Employment status** |  |  |  |  |  |
| Not employed(REF) | REF | REF | REF | REF | REF |
| Employed | 0.88 (0.49-1.57) ; 0.642 | 1.27 (0.8 - 2.03) ; 0.318 | 2.45 (1.17 - 5.17) ; 0.019 | 0.96 (0.58 - 1.58) ; 0.843 | 0.74 (0.39 - 1.41) ; 0.351 |

Abbreviations: OR, odds ratio; CI, confidence interval; FP, family planning

## DISCUSSION

We report the first data on *M. genitalium* in PNG, finding that pregnant women in PNG have one of the highest infections rates globally, with a striking absence of macrolide resistance despite resistance rates of up to 68%(18) in neighbouring regions such as Australia. The high prevalence of *M. genitalium* (12.5%) among pregnant women shown here equates to an estimated 13,000 prevalent cases (95% CI 10,342 to 15,823) of *M. genitalium* among women of reproductive age (15 to 49 years) in the province (Supplemental methods). Additionally, these are the first data on RTIs to be reported from the New Guinea Islands region of PNG in over twenty years and provides a more complete understanding of the burden of RTIs in pregnancy(26). This study indicates that more than one in two women (55.2%) have a treatable RTI (BV and or current STI) that causes harmful sexual and reproductive health outcomes and is not detected by current antenatal screening in PNG. Given that PNG has one of the largest populations among Pacific islands, these findings have major public health and regional significance.

There is currently no global *M. genitalium* surveillance, limiting our understanding of its epidemiology. High-income countries report rates of *M. genitalium* infection ranging from 0.3 to 3.3%(11, 13, 27) in the general population, with higher estimates in certain risk groups(28, 29). Fewer data are available from low- and middle-income countries (LMICs) but it appears prevalence may be higher ranging from 3% in the general population in Tanzania(13) to 8-9% in Honduras and South Africa(13, 30). The highest burden has been reported amongst those who sell sex, 16% in Kenya(31) and 26% in Uganda(32). Data specific to pregnancy remains limited despite the association with adverse pregnancy outcomes(28); in the United Kingdom and France prevalence among pregnant women is low (0.7-0.8%(14, 33)) with higher rates reported from Guinea-Bissau (6.2%)(34) and the Solomon Islands (11.9%)(15). Clearly more data on the burden of *M. genitalium* in pregnancy, and its consequences, are needed.

Recent data from the Solomon Islands examined the impact of mass drug administration (MDA) of 1g of azithromycin orally for the elimination of ocular *C. trachomatis* on *M. genitalium* and reported a pre-MDA *M. genitalium* prevalence of 11.9% (95% CI 8.3 – 16.6%; n=236) among women attending antenatal clinics. Post MDA *M. genitalium* prevalence remained high at 10.9% with no evidence of macrolide resistance in either pre- or post-MDA groups; however this is not surprising given only 5 of the 28 women positive for *M. genitalium* in the post-MDA group reported receiving azithromycin(15). The lack of macrolide resistance in *M. genitalium* infections in PNG found in this study was striking and warrants further exploration in other populations in PNG. While it may reflect a lack of exposure to macrolides in this population, macrolides are available and used widely in PNG, and their use has repeatedly been associated with an increase in macrolide resistant isolates in other settings.

The burden of curable STIs observed among pregnant women reported here is substantially greater than most settings included in the 2018 global estimates of curable STIs(3). The 32.1% observed prevalence of at least one current STI diagnosable by GeneXpert is lower than the 42.7% reported in a study of antenatal clinics from Eastern Highlands, Hela and Central Provinces of PNG in 2014(20), but similar to that reported in Madang Province in 2012 (33.7%)(35). *C. trachomatis* prevalence (19.1%) is consistent with recent reports from other provinces (Vallely 2016 (22.9%), Badman 2016 (20.0%)) and neighbouring Solomon Islands (20.3%)(36). These latter three studies were the highest reported rates from any study in the global prevalence estimates(3). Similarly for *N. gonorrhoea* in pregnancy, PNG and Solomon Islands have the highest reported rates globally (5.1% to 14.2%)(35–38), although two studies from South Africa also report very high rates at 10.1% in a primary care setting(39) and 6.4% among pregnant women(40). Regarding *T. vaginalis*, recent PNG estimates among pregnant women are generally higher than the 20.2% reported in this study (21.3-37.6%.(35, 37, 38, 41)).

Risk factors for STIs identified in this study (primigravida, employed, single/separated or having abnormal vaginal discharge at some point in the pregnancy up to ANC1) could have a number of explanations. Younger women in their first pregnancy may have had less interaction with reproductive health services, and employed women may be more mobile with an associated increase risk of STI acquisition through unprotected sex. Interestingly, no risk factors were identified for *M. genitalium*. Risk factors for STIs in pregnancy reported elsewhere in PNG include more than one lifetime sexual partner, level of education of the woman or her partner, rurality, previous miscarriage or stillbirth, and socioeconomic status(20, 35).

This study also provides important data regarding BV and VVC, with 58.3% of women having at least one of these infections. VVC is readily treatable(42) and can cause extreme discomfort in addition to increasing a woman’s risk of post-partum breast candidiasis with potential impact on breastfeeding. The prevalence reported here (39.4%) is higher than the only previous report from PNG in 1991 (23%)(43), comparisons with other LMICs are difficult as data is limited and not contemporary(42, 44). The prevalence of one in four women with BV reported here is higher than previous reports in PNG (17.6%)(41) but in keeping with recent global estimates of 23-29%(5). However, our results may underestimate the true burden of disease as diagnosis was limited to those with a Nugent’s score of seven to ten.

Syndromic management of RTIs is a pragmatic approach in the absence of definitive diagnosis. However, as has been reported in PNG and elsewhere(30, 37), we found this approach performed poorly, missing 78.2% of *M. genitalium* infections and three quarters of any RTI. Symptoms were associated with a higher odds of *T. vaginalis* infection; however, sensitivity remained low. It is clear that syndromic management is an inadequate tool to effectively treat RTIs, even for those infections more commonly associated with vaginal discharge. Improved access to affordable, accurate point-of-care diagnostics may be transformative. There is an urgent need for alternative approaches to more effectively detect and treat asymptomatic RTIs and reduce prevalence in a cost-effective and feasible manner in resource-limited settings.

The main limitation of this study is the facility-based recruitment of participants; results may not represent women who do not attend any antenatal clinic. Routinely collected provincial data for the study years estimated 73% to 85% of pregnant women attended clinic at least once during their pregnancy(45). A limitation of the syphilis data is that it represents lifetime exposure to *Treponema* species, and as such does not differentiate between active or latent infection, nor are we able to exclude exposure to yaws. Yaws is endemic in PNG(46), however prevalence estimates vary widely by region and precise estimates are not known for ENB. A population wide survey on an island in neighbouring New Ireland Province estimated a population prevalence of 1.8% for active yaws(47). Additionally, coverage with syphilis rapid tests was affected by supply interruptions.

This study provides the first data on *M. genitalium* prevalence and drug resistance markers in PNG, revealing a high burden of infection that is currently unrecognised, and provides valuable new data to understand the burden of this infection among pregnant women globally. STIs in pregnancy were alarmingly common, with 37.8% of pregnant women having at least one current STI. This study also highlights the high burden of bacterial vaginosis and VCC and clearly shows that current antenatal screening with syndromic management is inadequate in detecting reproductive tract infections. Such a high burden of disease with associated impacts on poor sexual and reproductive health demands urgent action towards ensuring access to affordable prevention, diagnostics and treatment for communities in PNG and similar settings. This will be crucial for achieving progress towards the sustainable development goals and improving reproductive health outcomes.

## NOTES

## Acknowledgements

The authors would like to extend our heartfelt thanks to the women and infants who participated in this study, as well as the families and communities who supported them to do so. Our special thanks to the National Department of Health, the East New Britain Provincial Administration led by Mr Wilson Matava, the Provincial Health Authority, Catholic Health Services and participating health facilities (Nonga General Hospital, St Mary’s Vunapope, Keravat rural hospital, Napapar health centre, Paparatava health centre) for enthusiastically facilitating our research team to work alongside them. Specific thanks to Mr Levi Mano and Mr Nicholas Larme, Dr Ako Yap, Mr Moses Bogandri, Mr Benedict Mode, Dr Pinip Wapi, Dr Felix Diaku, Dr Tanmay Bagade, Dr Delly Babona, Sr Placidia Nohan, Sr Theonila Wat and Sr Rebecca Penaia who have provided invaluable support and advice throughout the planning and implementation of this work in ENB. We gratefully acknowledge the dedication and contribution by our Burnet Institute Kokopo staff who worked tirelessly to implement this study, specifically we would like to thank: Dr Stenard Hiasihri, Essie Koniel, Pele Melepia, Hadlee Supsup, Dukduk Kabiu, Ruth Fidelis, Wilson Philip, Priscah Hezeri, Kerryanne Tokmun, Primrose Homiehombo, Rose Suruka, Benishar Kombut, Thalia Wat, Noelyne Taraba, Chris Sohenaloe, Dorish Palagat, Zoe Saulep, Elizabeth Walep, Lucy Au, Irene Daniels, Gabriella Kalimet-Tade, Noreen Tamtilik, Ellen Kavang, Wilson Kondo, Allan Tirang, Michael Palauva, Ioni Pidian, Teddy Wanahau, Eremas Amos, Bettie Matonge, Elice Adimain, Thelma Punion, Lucy Palom. Thank you to the invaluable project support from Burnet Institute Melbourne, especially: Kellie Woiwod, James Lawson, Lisa Davidson, Vivian Newton, Lisa Vitasovich and Rodney Stewart. We also thank our many collaborators, specifically Prof John Kaldor and Prof Rebecca Guy for advice on STIs. And for the vision, overall leadership and technical guidance to the HMHB program provided by Prof Michael Toole, Prof Margaret Hellard and Prof Caroline Homer.

## Authors contribution statement

Study design: led by JGB, BSC, FJIF, CM, MJLS with input from WP, SL, EK, PMS, LJR, AV, SGB, AJU. Data collection: MJLS, PM, EP, RF. Vaginal smears read by GF. Data analysis, interpretation and manuscript led by MJLS with input from all authors. All authors read and approved the final manuscript.

## Biographical Sketch of first author

**Dr Michelle Scoullar** is an international health specialist with extensive experience in maternal, newborn and child health, health system strengthening, and implementation research with practical clinical and public health program experience in remote Australia, Lao PDR and Papua New Guinea (PNG) and is a practicing medical doctor specialising in neonatology and paediatrics, with additional postgraduate qualifications in international health and obstetrics and gynaecology. Her primary research interests focus around health, infection and nutrition in pregnancy and subsequent impacts on neonatal and infant health, particularly birth weight and growth through infancy.

## REFERENCES

1. Newman L, Rowley J, Hoorn SV, Wijesooriya NS, Unemo M, Low N, et al. Global Estimates of the Prevalence and Incidence of Four Curable Sexually Transmitted Infections in 2012 Based on Systematic Review and Global Reporting. PloS one. 2015;10(12).

2. World Health Organization. Report on global sexually transmitted infection surveillance. 2018.

3. Rowley J, Hoorn SV, Korenromp E, Low N, Unemo M, Abu-Raddad LJ, et al. Chlamydia, gonorrhoea, trichomoniasis and syphilis: Global prevalence and incidence estimates, 2016. Bulletin of the World Health Organization. 2019;97(8):548–62, 62A-62P.

4. van de Wijgert JHHM, Jespers V. The global health impact of vaginal dysbiosis. Research in Microbiology. 2017;168(9-10):859–64.

5. Peebles K, Velloza J, Balkus JE, McClelland RS, Barnabas RV. High Global Burden and Costs of Bacterial Vaginosis: A Systematic Review and Meta-Analysis. Sexually Transmitted Diseases. 2019;46(5):304–11.

6. Cauchie M, Desmet S, Lagrou K. Candida and its dual lifestyle as a commensal and a pathogen. Research in Microbiology. 2017 2017/11/01/;168(9):802–10.

7. Donovan B. Sexually transmissible infections other than HIV. The Lancet. 2004;363(9408):545–56.

8. Arol OA, Over M, Manhard L, Holmes KK. Sexually Transmitted Infections. In: Dean T Jamison, Joel G Breman, Anthony R Measham, George Alleyne, Mariam Claeson, David B Evans, et al., editors. Disease control priorities in developing countries. 2nd ed. New York: Oxford University Press; 2006.

9. Thwaites A, Flanagan K, Datta S. Non-HIV sexually transmitted infections in pregnancy. Obstetrics, Gynaecology & Reproductive Medicine. 2019;29(6):151–7.

10. Martin DH, Manhart LE, Workowski KA. Mycoplasma genitalium From Basic Science to Public Health: Summary of the Results From a National Institute of Allergy and Infectious Disesases Technical Consultation and Consensus Recommendations for Future Research Priorities. The Journal of infectious diseases. 2017 Jul 15;216(suppl_2):S427–S30.

11. Lis R, Rowhani-Rahbar A, Manhart LE. Mycoplasma genitalium infection and female reproductive tract disease: a meta-analysis. Clinical infectious diseases: an official publication of the Infectious Diseases Society of America. 2015 Aug 1;61(3):418–26.

12. Sonnenberg P, Ison CA, Clifton S, Field N, Tanton C, Soldan K, et al. Epidemiology of Mycoplasma genitalium in British men and women aged 16-44 years: Evidence from the third National Survey of Sexual Attitudes and Lifestyles (Natsal-3). International Journal of Epidemiology. 2015;44(6):1982–94.

13. Baumann L, Cina M, Egli-Gany D, Goutaki M, Halbeisen FS, Lohrer G-R, et al. Prevalence ofMycoplasma genitaliumin different population groups: systematic review andmeta-analysis. Sexually transmitted infections. 2018;94(4):255–62.

14. Oakeshott P, Hay P, Taylor-Robinson D, Hay S, Dohn B, Kerry S, et al. Prevalence of Mycoplasma genitalium in early pregnancy and relationship between its presence and pregnancy outcome. BJOG: An International Journal of Obstetrics and Gynaecology. 2004;111(12):1464–7.

15. Harrison MA, Harding-Esch EM, Marks M, Pond MJ, Butcher R, Solomon AW, et al. Impact of mass drug administration of azithromycin for trachoma elimination on prevalence and azithromycin resistance of genital Mycoplasma genitalium infection. Sexually transmitted infections. 2019.

16. Lau A, Bradshaw CS, Lewis D, Fairley CK, Chen MY, Kong FY, et al. The Efficacy of Azithromycin for the Treatment of Genital Mycoplasma genitalium: A Systematic Review and Meta-analysis. Clinical infectious diseases: an official publication of the Infectious Diseases Society of America. 2015 Nov 1;61(9):1389–99.

17. Sweeney EL, Trembizki E, Bletchly C, Bradshaw CS, Menon A, Francis F, et al. Levels of mycoplasma genitalium antimicrobial resistance differ by both region and gender in the state of Queensland, Australia: Implications for treatment guidelines. Journal of Clinical Microbiology. 2019;57(3).

18. Read TRH, Fairley CK, Murray GL, Jensen JS, Danielewski J, Worthington K, et al. Outcomes of resistance-guided sequential treatment of mycoplasma genitalium infections: A prospective evaluation. Clinical Infectious Diseases. 2019;68(4):554–60.

19. The World Bank Group. Papua New Guinea. 2019; Available from: https://data.worldbank.org/country/papua-new-guinea.

20. Vallely LM, Toliman P, Ryan C, Rai G, Wapling J, Tomado C, et al. Prevalence and risk factors of Chlamydia trachomatis, Neisseria gonorrhoeae, Trichomonas vaginalis and other sexually transmissible infections among women attending antenatal clinics in three provinces in Papua New Guinea: a cross-sectional survey. Sex Health. 2016 Oct;13(5):420–7.

21. Mola G, Amoa A, Bagita M, Augerea L, Geita L, O’Connor M. Manual of Standard Managements in Obstetrics and Gynaecology for Doctors, HEOs and Nurses in Papua New Guinea. 7th ed. Glen DL Mola, editor. Port Moresby, Papua New Guinea: World Health Organization; 2016.

22. National Department of Health. Standard Treatment Guidelines for common illness of adults in Papua New Guinea. 6th ed. Manning L, editor. Port Moresby, Papua New Guinea: World Health Organization; 2012.

23. Tabrizi SN, Su J, Bradshaw CS, Fairley CK, Walker S, Tan LY, et al. Prospective evaluation of ResistancePlus MG, a new multiplex quantitative PCR assay for detection of Mycoplasma genitalium and macrolide resistance. Journal of Clinical Microbiology. 2017;55(6):1915–9.

24. Su JP, Tan LY, Garland SM, Tabrizi SN, Mokany E, Walker S, et al. Evaluation of the SpeeDx ResistancePlus MG diagnostic test for mycoplasma genitalium on the applied biosystems 7500 fast quantitative PCR platform. Journal of Clinical Microbiology. 2018;56(1).

25. Pitt R, Cole MJ, Fifer H, Woodford N. Evaluation of the Mycoplasma genitalium Resistance Plus kit for the detection of M. genitalium and mutations associated with macrolide resistance. Sexually transmitted infections. 2018;94(8):565–7.

26. Vallely A, Page A, Dias S, Siba P, Lupiwa T, Law G, et al. The prevalence of sexually transmitted infections in Papua New Guinea: a systematic review and meta-analysis. PloS one. 2010;5(12):e15586.

27. Jensen JS, Cusini M, Gomberg M, Moi H. 2016 European guideline on Mycoplasma genitalium infections. J Eur Acad Dermatol Venereol. 2016 Oct;30(10):1650–6.

28. Donders GGG, Ruban K, Bellen G, Petricevic L. Mycoplasma/Ureaplasma infection in pregnancy: To screen or not to screen. Journal of Perinatal Medicine. 2017;45(5):505–15.

29. Deborde M, Pereyre S, Puges M, Bébéar C, Desclaux A, Hessamfar M, et al. High prevalence of Mycoplasma genitalium infection and macrolide resistance in patients enrolled in HIV pre-exposure prophylaxis program. Medecine et Maladies Infectieuses. 2019;49(5):347–9.

30. Hoffman CM, Mbambazela N, Sithole P, Morré SA, Dubbink JH, Railton J, et al. Provision of Sexually Transmitted Infection Services in a Mobile Clinic Reveals High Unmet Need in Remote Areas of South Africa: A Cross-sectional Study. Sexually Transmitted Diseases. 2019;46(3):206–12.

31. Cohen CR, Nosek M, Meier A, Astete SG, Iverson-Cabral S, Mugo NR, et al. Mycoplasma genitalium infection and persistence in a cohort of female sex workers in Nairobi, Kenya. Sex Transm Dis. 2007 May;34(5):274–9.

32. Vandepitte J, Muller E, Bukenya J, Nakubulwa S, Kyakuwa N, Buve A, et al. Prevalence and correlates of Mycoplasma genitalium infection among female sex workers in Kampala, Uganda. The Journal of infectious diseases. 2012 Jan 15;205(2):289–96.

33. Peuchant O, Le Roy C, Desveaux C, Paris A, Asselineau J, Maldonado C, et al. Screening for Chlamydia trachomatis, Neisseria gonorrhoeae, and Mycoplasma genitalium should it be integrated into routine pregnancy care in French young pregnant women? Diagn Microbiol Infect Dis. 2015 May;82(1):14–9.

34. Labbe AC, Frost E, Deslandes S, Mendonca AP, Alves AC, Pepin J. Mycoplasma genitalium is not associated with adverse outcomes of pregnancy in Guinea-Bissau. Sexually transmitted infections. 2002 Aug;78(4):289–91.

35. Wangnapi RA, Soso S, Unger HW, Sawera C, Ome M, Umbers AJ, et al. Prevalence and risk factors for Chlamydia trachomatis, Neisseria gonorrhoeae and Trichomonas vaginalis infection in pregnant women in Papua New Guinea. Sexually transmitted infections. 2015 May;91(3):194–200.

36. Marks M, Kako H, Butcher R, Lauri B, Puiahi E, Pitakaka R, et al. Prevalence of sexually transmitted infections in female clinic attendees in Honiara, Solomon Islands. BMJ open. 2015;5(4):e007276.

37. Vallely LM, Toliman P, Ryan C, Rai G, Wapling J, Gabuzzi J, et al. Performance of syndromic management for the detection and treatment of genital Chlamydia trachomatis, Neisseria gonorrhoeae and Trichomonas vaginalis among women attending antenatal, well woman and sexual health clinics in Papua New Guinea: a cross-sectional study. BMJ open. 2017 Dec 29;7(12):e018630.

38. Unger HW, Ome-Kaius M, Wangnapi RA, Umbers AJ, Hanieh S, Suen CSNLW, et al. Sulphadoxine-pyrimethamine plus azithromycin for the prevention of low birthweight in Papua New Guinea: A randomised controlled trial. BMC Medicine. 2015;13(1).

39. Peters RPH, Dubbink JH, Van Der Eem L, Verweij SP, Bos MLA, Ouburg S, et al. Cross-sectional study of genital, rectal, and pharyngeal chlamydia and gonorrhea in women in rural South Africa. Sexually Transmitted Diseases. 2014;41(9):564–9.

40. Moodley D, Moodley P, Sebitloane M, Soowamber D, McNaughton-Reyes HL, Groves AK, et al. High prevalence and incidence of asymptomatic sexually transmitted infections during pregnancy and postdelivery in KwaZulu Natal, South Africa. Sexually Transmitted Diseases. 2015;42(1):43–7.

41. Badman SG, Vallely LM, Toliman P, Kariwiga G, Lote B, Pomat W, et al. A novel point-of-care testing strategy for sexually transmitted infections among pregnant women in high-burden settings: results of a feasibility study in Papua New Guinea. BMC Infectious Diseases. 2016;16(1).

42. Pappas PG, Kauffman CA, Andes D, Benjamin Jr DK, Calandra TF, Edwards Jr JE, et al. Clinical practice guidelines for the management of candidiasis: 2009 Update by the Infectious Diseases Society of America. Clinical Infectious Diseases. 2009;48(5):503–35.

43. Klufio CA, Amoa AB, Delamare O, Hombhanje M, Kariwiga G, Igo J. Prevalence of vaginal infections with bacterial vaginosis, Trichomonas vaginalis and Candida albicans among pregnant women at the Port Moresby General Hospital Antenatal Clinic. Papua and New Guinea medical journal. 1995;38(3):163–71.

44. Sobel JD. Vulvovaginal candidosis. Lancet. 2007;369(9577):1961–71.

45. Papua New Guinea National Department of Health. Sector Performance Annual Review. 2018.

46. Mitjà O, Marks M, Konan DJP, Ayelo G, Gonzalez-Beiras C, Boua B, et al. Global epidemiology of yaws: a systematic review. The Lancet Global Health. 2015;3(6):e324–e31.

47. Mitjà O, Godornes C, Houinei W, Kapa A, Paru R, Abel H, et al. Re-emergence of yaws after single mass azithromycin treatment followed by targeted treatment: a longitudinal study. The Lancet. 2018;391(10130):1599–607.

