## Supplemental Material for "High burden of *Mycoplasma genitalium* and other reproductive tract infections among pregnant women in Papua New Guinea"

**Supplemental Methods:**

**Study procedures:**

*Routine antenatal care provided in line with PNG national guidelines*

Routine prevention and treatment care includes intermittent preventive treatment in pregnancy (IPTp) with sulphadoxine-pyrimethamine (SP) for malaria, iron and folate supplementation, voluntary counselling and testing for human immunodeficiency virus (HIV) and syphilis screening at point-of-care (POC) using Alere Determine™ HIV-1/2 and Alere Determine™ Syphilis TP. As per national syndromic management guidelines, pregnant women reporting current abnormal vaginal discharge receive: an intravaginal nystatin pessary (100,000 units, twice daily for 7 days) or a clotrimazole pessary once followed by intravaginal clotrimazole cream for 7 days; amoxycillin 2g orally (PO); probenecid 1g PO; augmentin 2 tablets PO, and Azithromycin 1g PO. Those with genital ulcer syndrome (current genital sores or ulcers) were treated for syphilis and if lesion not resolved after 1 week, treated for Donovanosis.

*Sample collection*

Two self-collected vaginal swabs were provided by participants, one GeneXpert® vaginal/endocervical swab and one Copan flocked swab. The GeneXpert swab was placed directly into its transport medium, and the Copan swab was placed in Copan Universal Transport Medium (UTM-RT) after first being used to prepare a vaginal smear on a slide for microscopy; both specimens were stored in a chilled esky for the remainder of the clinic. Urine samples were collected in a sterile container which was placed on ice and later stored at -20 degrees Celsius. On the same day as collection, specimens were transported to the Burnet Institute laboratory in Kokopo for processing, and storage. Copan UTM-RT tubes were stored at -20°C for later shipment, Xpert® swabs were tested for *C.trachomatis, N.gonorrhoea and T. vaginalis* and the vaginal smears were air dried for 24 hours then shipped to Burnet Institute Melbourne, Australia in batches. In cases where testing was delayed, specimens were stored at 2-7 degrees Celsius or at -20 degrees if an extended delay was anticipated. Testing usually occurred within seven days of specimen collection with results communicated back to participants and treatment advised for themselves and their partner. Remaining vaginal and urine specimens were stored at -20°C and later shipped frozen to the Burnet Institute in Melbourne, Australia.

**Laboratory methods**

The GeneXpert platform (Cepheid, Sunnyvale CA, USA) was used on available vaginal specimens or urine at Burnet Institute Kokopo (within 1 hour of all participating health facilities) for *C.trachomatis, N.gonorrhoea and T. vaginalis*. Lifetime exposure to *T. pallidum* (syphilis) was determined to be positive if women tested positive by Alere Determine™ Syphilis TP.

Shipped vaginal specimens were stored at Burnet Institute Melbourne at -20 or -80°C until all samples had been received. After thawing, genomic DNA was extracted using the QIAamp® BiOstic® Bacteremia DNA (QIAGEN) kit. To optimise this protocol for vaginal specimens rather than cultured blood the total volume of sample required was adjusted to 450μl. Extracted DNA was tested for *M. genitalium* using the *ResistancePlus*® MG kit (SpeeDx, Sydney, Australia) which uses the PlexPCR® and PlexPrime® technologies (SpeeDx) for concurrent amplification of *M. genitalium* and detection of five *M. genitalium* point mutations (A2058G, A2058C, A2058T, A2059G, and A2059C) within the macrolide resistance-determining region (MRDR) of the 23S rRNA gene, each of these point mutations are independently associated with macrolide resistance.(17) The *ResistancePlus*® MG kit is commercially available, TGA approved, and has a sensitivity and specificity of 98% and 100% for *M. genitalium* and 100% and 96% for macrolide resistance mutations.(23-25) Vaginal smears were read by an experienced microscopist at the Melbourne Sexual Health Centre. BV diagnosis was based on Nugent scoring (BV, Nugent score 7 to 10) and presumptive diagnosis of candidiasis was based on the observation of pseudohyphae and/or budding yeasts*.* Because the diagnosis of *M. genitalium,* BV and VVC was on stored samples many months after collection it was not possible to specifically treat those women who tested positive beyond any syndromic management they would have received as part of national guidelines.

**Exposures and outcomes**

We determined predictors of RTI at first antenatal clinic visit. Exposures of interest included clinic details (enrolment clinic, clinic location as rural or urban); participant characteristics at enrolment including age group (categorised as 16 – 24; 25 – 34 ; and 35 years or older), gravidity (primigravida or multigravida), marital status (partnered or single/separated), history of ever having used a modern contraceptive (WHO definition: oral contraceptive pills, implants, injectables, female sterilisation, male sterilisation, intra-uterine devices, diaphragm, emergency contraception, male and female condoms)(26), vaginal discharge, religion, employment status, education, smoking, alcohol use, mid upper arm circumference (MUAC; categorised as ≤23cm, >23cm), self-reported history of STI, presence of nitrites on urine dipstick (as a proxy for urinary tract infection), previous pregnancy outcome (miscarriage, livebirth or stillbirth), haemoglobin level in g/dl and WHO anaemia classification for pregnant women at sea level (mild anaemia 10-10.9g/dl; moderate 7.0-9.9g/dl; severe <7.0g/dl)(27); household characteristics including partner education, polygamy, time travelled to clinic, household monthly expenditure. Outcomes measured were *M. genitalium, C.trachomatis, N.gonorrhoea, T. vaginalis, T. pallidum* (syphilis), bacterial vaginosis and vaginal candidiasis.

**Data management**

Questionnaire responses were entered by the research officer directly into an electronic tablet (e-tablet) using a study specific questionnaire on the platform Mobile Data Studio 7.3 (MDS, CreativityCorp Pty Ltd, Perth, Australia). After each clinic, questionnaires were uploaded into a central computer, verified for completeness and exported from MDS to Microsoft® Office Excel. GeneXpert laboratory results were uploaded by the study data manager on a weekly basis and results checked for accuracy and exported. Laboratory results generated in Melbourne for *M. genitalium*, BV and VVC were exported to Microsoft Excel. All Excel files were provided to the first author and uploaded into STATA 15.0 (Stata Inc., College Station, TX, USA). The databases were verified and cleaned using STATA 15.0 and merged into one database for analysis. All data was stored on password protected computers accessible only to research personnel.

**Ethical considerations and informed consent procedures**

All participants provided individual written informed consent to participate. Ethical approval was provided from the Medical Research Advisory Committee of the PNG National Department of Health (No. 14.27), the PNG Institute of Medical Research (PNG IMR) Institutional Review Board (No. 1114) and the Human Research Ethics Committee of the Alfred Hospital (No. 348/14) in Australia. Provincial approval was obtained from the East New Britain Province (ENBP) executive committee, participating facilities and a series of community engagement meetings provided broader community support and assent for the study.

**Calculation for number of *M. genitalium* prevalent cases in East New Britain Province.**

Recruitment occurred over a period of 26 months and 3 weeks (116.14 weeks, 813 days) from 16^th^ March 2015 through to 6^th^ June 2017. Midpoint for recruitment is therefore 13.3 months (58.1 weeks, 406 days) after recruitment began, so the 25^th^ April 2016.

The population of East New Britain at the 2011 census was 328,369 (taken the 10^th^ July 2011), representing an annual growth rate of 3.6% since the previous census.

PNG total number of women in 2011 3,497,244, of which 1,932,533 (55.3%) were women of reproductive age (15 to 49 years). ENB Province had a total of 159,609 women in 2011, therefore the number of women of reproductive age (15 – 49 years) in ENB at 2011 Census = 159,609 * 0.553 = 88,263.8 women.

Years between 10^th^ July 2011 census and HMHB recruitment mid-point = 4 years, 9 months and 2 weeks or 4.769 years.

Calculation used to estimate number of women aged 15 – 49 years in ENB at study mid-point:

2011 census number	+	Population growth since census

88,263.8	+	(88,263.8 * 4.769 * 0.036)	= 15,153.478

= 103,417.278

Study estimated population prevalence for *M.genitalium* of 12.5% (95% CI 10.0-15.3). Therefore estimated number of prevalence cases of *M.genitalium* for 2016 = 103,417.278*0.125 = 12,927.159 (95%CI 10,341.728 – 15,822.843)

**SUPPLEMENTAL TABLE 1:** Socio-demographic characteristics and obstetric history of women who received RTI testing

|  |  |  | **Total at ANC1 n (%)** | Missing data MG | **MG test done n (%)** | Missing data MG | **CT/NG test done n (%)** | Missing data CT/NG | **TV test done n (%)** | Missing data TV | **Syphilis test done n (%)** | Missing data Syphilis | **BV and VVC test done n (%)** | Missing data BV and VVC |
| --- | --- | --- | --- | --- | --- | --- | --- | --- | --- | --- | --- | --- | --- | --- |
|  |  |  | N=699 unless specified | | n=625 |  | n=641 |  | n=581 |  | n=437 |  | n=503 |  |
| **Sociodemographic details for enrolled women** | | |  |  |  |  |  |  |  |  |  |  |  |  |
|  | Enrolment Clinic | |  |  |  |  |  |  |  |  |  |  |  |  |
|  |  | Vunapope | 184 (26.3) |  | 157 (25.1) |  | 169 (26.4) |  | 164 (28.2) |  | 89 (20.4) |  | 145 (28.8) |  |
|  |  | Nonga | 83 (11.9) |  | 81 (13.0) |  | 73 (11.4) |  | 64 (11.0) |  | 68 (15.6) |  | 69 (13.7) |  |
|  |  | Keravat | 125 (17.9) |  | 114 (18.2) |  | 118 (18.4) |  | 90 (15.5) |  | 86 (19.7) |  | 101 (20.1) |  |
|  |  | Napapar | 158 (22.6) |  | 136 (21.8) |  | 141 (22.0) |  | 131 (22.5) |  | 101 (23.1) |  | 115 (22.9) |  |
|  |  | Paparatava | 149 (21.3) |  | 137 (21.9) |  | 140 (21.8) |  | 132 (22.7) |  | 93 (21.3) |  | 73 (14.5) |  |
|  | Clinic administration | |  |  |  |  |  |  |  |  |  |  |  |  |
|  |  | Government | 208 (29.8) |  | 195 (31.2) |  | 191 (29.8) |  | 154 (26.5) |  | 154 (35.2) |  | 170 (33.8) |  |
|  |  | Catholic Health | 491 (70.2) |  | 430 (68.8) |  | 450 (70.2) |  | 427 (73.5) |  | 283 (64.8) |  | 333 (66.2) |  |
|  | Location | |  |  |  |  |  |  |  |  |  |  |  |  |
|  |  | Urban | 342 (48.9) |  | 293 (46.9) |  | 310 (48.4) |  | 295 (50.8) |  | 190 (43.5) |  | 260 (51.7) |  |
|  |  | Rural | 357 (51.1) |  | 332 (53.1) |  | 331 (51.6) |  | 286 (49.2) |  | 247 (56.5) |  | 243 (48.3) |  |
|  | Age, years | |  |  |  |  |  |  |  |  |  |  |  |  |
|  |  | mean [SD], range | 26.8 [5.6], 16-49 | 9 | 26.8 [5.5], 16-49 | 9 | 26.8 [5.6], 16-49 | 7 | 26.9 [5.6], 17-49 | 7 | 26.7 [5.6], 17-49 | 7 | 26.7 [5.6], 17-49 | 7 |
|  | Highest level of education completed | |  | 1 |  | 1 |  | 1 |  | 1 |  |  |  | 1 |
|  |  | Primary (Grade 8 or less) | 325 (46.6) |  | 302 (48.4) |  | 301 (47.0) |  | 274 (47.2) |  | 204 (46.7) |  | 229 (45.6) |  |
|  |  | High school (grade 9,10) | 177 (25.4) |  | 151 (24.2) |  | 160 (25.0) |  | 138 (23.8) |  | 112 (25.6) |  | 130 (25.9) |  |
|  |  | Secondary / Vocational / Tertiary | 196 (28.1) |  | 171 (27.4) |  | 179 (28.0) |  | 168 (29.0) |  | 121 (27.7) |  | 143 (28.5) |  |
|  | Employment status | |  |  |  |  |  |  |  |  |  |  |  |  |
|  |  | Not employed | 531 (76.0) |  | 475 (76.0) |  | 488 (76.1) |  | 443 (76.2) |  | 339 (77.6) |  | 373 (74.2) |  |
|  |  | Employed in paid work or student | 168 (24.0) |  | 150 (24.0) |  | 153 (23.9) |  | 138 (23.8) |  | 98 (22.4) |  | 130 (25.8) |  |
|  | Province of birth | |  |  |  |  |  |  |  |  |  |  |  |  |
|  |  | East New Britain | 578 (82.7) |  | 518 (82.9) |  | 529 (82.5) |  | 479 (82.4) |  | 362 (82.8) |  | 411 (81.7) |  |
|  |  | Other Province | 121 (17.3) |  | 107 (17.1) |  | 112 (17.5) |  | 102 (17.6) |  | 75 (17.2) |  | 92 (18.3) |  |
|  | Religion | |  | 1 |  | 1 |  | 1 |  | 1 |  | 1 |  | 1 |
|  |  | Catholic | 345 (49.4) |  | 302 (48.4) |  | 317 (49.5) |  | 281 (48.4) |  | 217 (49.8) |  | 234 (46.6) |  |
|  |  | Other | 353 (50.6) |  | 322 (51.6) |  | 323 (50.5) |  | 299 (51.6) |  | 219 (50.2) |  | 268 (53.4) |  |
|  | Marital status | |  | 2 |  | 2 |  | 2 |  | 2 |  |  |  | 2 |
|  |  | Married or cohabiting | 663 (95.1) |  | 593 (95.2) |  | 608 (95.1) |  | 550 (95.0) |  | 418 (95.7) |  | 477 (95.2) |  |
|  |  | Single, seperated or widowed | 34 ( 4.9) |  | 30 ( 4.8) |  | 31 ( 4.9) |  | 29 ( 5.0) |  | 19 ( 4.3) |  | 24 ( 4.8) |  |
|  | Poligamy | |  | 37 |  | 33 |  | 34 |  | 31 |  | 18 |  | 28 |
|  |  | One wife | 583 (88.1) |  | 523 (88.3) |  | 531 (87.5) |  | 483 (87.8) |  | 369 (88.1) |  | 416 (87.6) |  |
|  |  | More than one wife | 79 (11.9) |  | 69 (11.7) |  | 76 (12.5) |  | 67 (12.2) |  | 50 (11.9) |  | 59 (12.4) |  |
|  | Household monthly expenditure in Kina | | 150 {50-300} | 36 | 150 {50-300} | 35 | 150 {50-300} | 30 | 150 {60-300} | 32 | 150 {60-300} | 17 | 150 {99-300} | 24 |
|  | Cost of ANC in Kina | | 4 {2-20} | 17 | 4 {2-20} | 16 | 4 {2-20} | 15 | 4 {2-20} | 14 | 4 {2-20} | 12 | 5 {2-20} | 11 |
| **Family Planning** | | |  |  |  |  |  |  |  |  |  |  |  |  |
|  | Method used | |  | 9 |  | 9 |  | 9 |  | 9 |  | 4 |  | 3 |
|  |  | Never used modern FP method | 569 (82.5) |  | 510 (82.8) |  | 513 (81.2) |  | 469 (82.0) |  | 361 (83.4) |  | 411 (82.2) |  |
|  |  | Has used modern FP method | 121 (17.5) |  | 106 (17.2) |  | 119 (18.8) |  | 103 (18.0) |  | 72 (16.6) |  | 89 (17.8) |  |
| **Maternal Health Parameters at 1st Antenatal Clinic** | | | |  |  |  |  |  |  |  |  |  |  |  |
|  | Gravidity | |  |  |  |  |  |  |  |  |  |  |  |  |
|  |  | Primigravidae | 177 (25.3) |  | 159 (25.4) |  | 166 (25.9) |  | 145 (25.0) |  | 111 (25.4) |  | 125 (24.9) |  |
|  |  | Multigravidae (2-4) | 384 (54.9) |  | 343 (54.9) |  | 343 (53.5) |  | 318 (54.7) |  | 243 (55.6) |  | 284 (56.5) |  |
|  |  | Grandmulti (≥5) | 138 (19.7) |  | 123 (19.7) |  | 132 (20.6) |  | 118 (20.3) |  | 83 (19.0) |  | 94 (18.7) |  |
|  | Abnormal vaginal discharge | |  |  |  |  |  |  |  |  |  |  |  |  |
|  |  | Abnormal discharge at any time in preg | 563 (80.7) |  | 507 (81.1) |  | 512 (79.9) |  | 466 (80.2) |  | 358 (81.9) |  | 401 (79.7) |  |
|  |  | Current symptoms | 135 (19.3) |  | 118 (18.9) |  | 129 (20.1) |  | 115 (19.8) |  | 79 (18.1) |  | 102 (20.3) |  |
|  | Smoking | |  | 1 |  | 1 |  | 1 |  | 1 |  | 1 |  | 1 |
|  |  | Never smoked | 427 (61.3) |  | 384 (61.5) |  | 387 (60.5) |  | 352 (60.7) |  | 268 (61.5) |  | 305 (60.8) |  |
|  |  | Stopped when pregnant | 241 (34.6) |  | 217 (34.8) |  | 226 (35.3) |  | 204 (35.2) |  | 151 (34.6) |  | 174 (34.7) |  |
|  |  | Current smoker | 29 ( 4.2) |  | 23 ( 3.7) |  | 27 ( 4.2) |  | 24 ( 4.1) |  | 17 ( 3.9) |  | 23 ( 4.6) |  |
| **Previous pregnancy outcomes** | | |  |  |  |  |  |  |  |  |  |  |  |  |
|  | Age at first pregnancy | | 21 {19-24} | 7 | 21 {19-24} | 7 | 21 {19-24} | 5 | 21 {19-24} | 5 | 21 {19-24} | 5 | 21 {19-23} | 5 |
|  | History pregnancy loss (in multiparous) | |  |  | n=466 |  | n=475 |  | n=436 |  | n=326 |  | n=378 |  |
|  |  | History of miscarriage | 46 (8.8) |  | 44 (9.4) |  | 43 (9.1) |  | 39 (8.9) |  | 29 (8.9) |  | 34 (9.0) |  |
|  |  | History of abortion | 1 (0.2) |  | 1 (0.2) |  | 1 (0.2) |  | 1 (0.2) |  | 0 (0.0) |  | 0 (0.0) |  |
|  |  | History of stillbirth | 16 (3.1) |  | 14 (3.0) |  | 16 (3.4) |  | 14 (3.2) |  | 5 (1.5) |  | 10 (2.6) |  |
| **Partner details** | | |  |  |  |  |  |  |  |  |  |  |  |  |
|  | Partner's employment status | |  | 18 |  | 18 |  | 17 |  | 14 |  | 10 |  | 12 |
|  |  | Not employed | 269 (39.6) |  | 240 (39.5) |  | 249 (39.9) |  | 229 (40.4) |  | 165 (38.6) |  | 181 (36.9) |  |
|  |  | Employed in paid work | 411 (60.4) |  | 367 (60.5) |  | 375 (60.1) |  | 338 (59.6) |  | 262 (61.4) |  | 310 (63.1) |  |
|  | Partner attending ANC1 | |  | 3 |  | 3 |  | 3 |  | 2 |  | 2 |  | 4 |
|  |  | No | 571 (82.3) |  | 518 (83.3) |  | 529 (82.9) |  | 477 (82.4) |  | 363 (83.4) |  | 418 (83.8) |  |
|  |  | Yes | 123 (17.7) |  | 104 (16.7) |  | 109 (17.1) |  | 102 (17.6) |  | 72 (16.6) |  | 81 (16.2) |  |
| **data are mean [SD], range; or median {IQR}, range; or n (%)** | | | | | | | | | | | | | | |

**SUPPLEMENTAL TABLE 2:** Relationship between Active STIs

|  | **Total (N)** | **Mono-infection n (%)** | **Co-infection n (%)** | ***M. genitalium*** | ***C. trachomatis*** | ***N. gonorrhoeae*** | ***T. vaginalis*** |
| --- | --- | --- | --- | --- | --- | --- | --- |
| ***Mycoplasma genitalium*** | 78 | 32 (41.0) | 28 (35.9) | - | 20 | 6 | 13 |
| ***Chlamydia trachomatis*** | 122 | 36 (29.5) | 66 (54.1) | 20 | - | 25 | 36 |
| ***Neisseria gonorrhoeae*** | 35 | 4 (11.4) | 28 (80.0) | 6 | 25 | - | 8 |
| ***Trichomonas vaginalis*** | 117 | 56 (47.8) | 44 (37.6) | 13 | 36 | 8 | - |
| **NB:** Totals do not equal sum of co-infections as some women had multiple infections and not all women received testing for all infections | | | | | | | |

**SUPPLEMENTAL TABLE 3**: Relationship between BV and active STIs

|  | **Total (N)** | **Mono-infection n (%)** | **Co-infection n (%)** | ***M. genitalium*** | ***C. trachomatis*** | ***N. gonorrhoeae*** | ***T. vaginalis*** |
| --- | --- | --- | --- | --- | --- | --- | --- |
| **Bacterial Vaginosis** | 129 | 59 (45.7) | 48 (37.2) | 15 | 29 | 10 | 16 |
| **NB:** Total does not equal sum of mono- and co-infections as some women had multiple infections and not all women received testing for all infections | | | | | | | |

| **Supplemental Table 4.** Sensitivity and specificity of clinical symptoms as a marker of Reproductive Tract Infections (RTIs) | | | | | | | | |
| --- | --- | --- | --- | --- | --- | --- | --- | --- |
|  |  | **Question as per syndromic management** | |  |  | **Alternative question** | |  |
|  |  | Do you currently have any abnormal vaginal discharge? | | |  | Have you experienced any abnormal vaginal discharge earlier in the pregnancy or now? | | |
| **Reproductive Tract Infection** |  | **p-value^b^** | **Sensitivity / Specificity (95%CI)** | **NPV / PPV (95%CI)** |  | **p-value^b^** | **Sensitivity / Specificity (95%CI)** | **NPV / PPV (95%CI)** |
| *Mycoplasma genitalium* |  | 0.628 | 15.4 (8.3-25.4) / 86.7 (83.5-89.4) | 14.2 (7.6-23.4) / 87.8 (84.7-90.5) |  | 0.482 | 21.8 (13.3-32.6) / 81.6 (78.1 - 84.8) | 14.5 (8.7-22.1) / 88 (84.9 - 90.7) |
| *Chlamydia trachomatis* |  | 0.075 | 19.7 (13.1-27.9) / 86.7 (83.5-89.5) | 25.9 (17.3-36) / 82.1 (78.6-85.2) |  | 0.063 | 26.3 (18.7-35) / 81.3 (77.7 - 84.6) | 24.9 (17.7-33.2) / 82.4 (78.9 - 85.6) |
| *Neisseria gonorrhoeae* |  | 0.347 | 20 (8.5-37) / 85.8 (82.8-88.5) | 7.6 (3.1-14.9) / 94.9 (92.7-96.6) |  | 0.399 | 25.8 (12.5-43.3) / 80.2 (76.8 - 83.3) | 7 (3.3-12.9) / 95 (92.7 - 96.7) |
| *Trichomonas vaginalis* |  | 0.047 | 19.7 (12.9-28.1) / 87.5 (84.2-90.4) | 28.4 (19-39.6) / 81.2 (77.5-84.6) |  | 0.005 | 29.1 (21.1-38.2) / 82.6 (78.8 - 85.9) | 29.6 (21.5-38.8) / 82.2 (78.5 - 85.6) |
| Bacterial Vaginosis |  | 0.777 | 15.6 (9.8-23) / 85.6 (81.6-89) | 27.1 (17.4-38.7) / 74.6 (70.2-78.6) |  | 0.640 | 21.8 (15-29.9) / 80.3 (75.9 - 84.2) | 27.5 (19.1-37.2) / 74.9 (70.3 - 79) |
| Vulvovaginal Candidiasis |  | 0.023 | 19.2 (14-25.4) / 88.2 (84-91.6) | 51.4 (39.5-63.2) / 62.7 (57.9-67.3) |  | 0.007 | 26.3 (20.3-33) / 83.7 (79 - 87.6) | 51 (40.9-61.1) / 63.6 (58.7 - 68.4) |
| **Multiple Infections** |  |  |  |  |  |  |  |  |
| At least 1 active RTI^c^ |  | 0.009 | 17.7 (13.4-22.8) / 94.1 (86.7-98.1) | 90.6 (79.4-96.9) / 26.1 (21.3-31.5) |  | 0.005 | 25.0 (20-30.6) / 89.5 (80.9 - 95.1) | 88.4 (79-94.6) / 27.2 (22.1 - 32.8) |
| At least 1 active STI^d^ |  | 0.224 | 15.9 (10.9-22) / 88.1 (83.9-91.5) | 44.7 (32.3-57.5) / 63.3 (58.5-67.9) |  | 0.146 | 23.0 (17.1-29.8) / 82.5 (77.7 - 86.6) | 44.3 (34.1-54.8) / 63.9 (58.9 - 68.7) |
| At least 1 cervical infection^e^ |  | 0.793 | 14.5 (9.5-20.8) / 86.4 (82.7-89.6) | 30 (20.3-41.3) / 71.5 (67.3-75.4) |  | 0.846 | 19.9 (14.1-26.8) / 80.9 (76.7 - 84.6) | 29.5 (21.3-38.9) / 71.5 (67.2 - 75.6) |
| At least 1 vaginal infection^f^ |  | 0.001 | 18.9 (14.5-24) / 93.8 (88-97.3) | 86.9 (75.8-94.2) / 34.3 (29.4-39.6) |  | <0.001 | 26.4 (21.3-31.9) / 90.7 (84.2 - 95.1) | 86.1 (76.9-92.6) / 36 (30.7 - 41.5) |
| At least 1 GeneXpert diagnosed infection^g^ |  | 0.019 | 19.5 (13.9-26.1) / 88.2 (84.4-91.3) | 43.6 (32.4-55.3) / 69.9 (65.5-74) |  | 0.005 | 27.5 (21-34.7) / 83.1 (78.9 - 86.7) | 43.3 (33.9-53) / 70.9 (66.3 - 75.1) |
| Any 2 active STIs |  | 0.030 | 22.7 (13.8-33.8) / 86.7 (83.7-89.4) | 17.9 (10.8-27.1) / 89.8 (87-92.2) |  | 0.028 | 29.4 (19.4-41) / 81.4 (78.1 - 84.5) | 16.8 (10.9-24.4) / 90 (87.2 - 92.5) |
| ^a^: symptoms refer to abnormal vaginal discharge only (not genital warts/ulcers/sores)  ^b^ p-value derived from Pearson's χ^2^ test  ^c^ active RTI includes at least 1 of MG, CT, NG, TV, BC or VVC (syphilis not included)  ^d^ active STI includes at least 1 of MG, CT, NG, TV (syphilis not included)  ^e^ cervical infections includes at least 1 of MG, CT or NG  ^f^ vaginal infections includes at least 1 of BV, TV or VVC  g GeneXpert diagnosed infections include any 1 of CT, NG or TV  PPV: Positive predictive value; NPV: Negative predictive value  MG (Mycoplasma genitalium), CT (Chlamydia trachomatis), NG (Neisseria gonorrhoea), TV (Trichomonas vaginalis), VVC (vulvovaginal candidiasis) | | | | | | | | |
